# A modelling assessment for the impact of control measures on highly pathogenic avian influenza transmission in poultry in Great Britain

**DOI:** 10.1101/2025.04.24.650264

**Authors:** Christopher N Davis, Edward M Hill, Chris P Jewell, Kristyna Rysava, Robin N Thompson, Michael J Tildesley

## Abstract

Since 2020, large-scale outbreaks of highly pathogenic avian influenza (HPAI) H5N1 in Great Britain have resulted in substantial poultry mortality and economic losses. Alongside the costs, the risk of circulation leading to a viral reassortment that causes zoonotic spillover raises additional concerns. However, the precise mechanisms driving transmission between poultry premises and the impact of potential control measures in Great Britain, such as vaccination, are not fully understood. We have developed a spatial transmission model for the spread of HPAI in poultry premises calibrated to infected premises data for the 2022–23 season using Markov chain Monte Carlo. Our results indicate that enhanced biosecurity measures and/or vaccination of the premises surrounding an identified infected premises can substantially reduce the overall number of infected premises. Our findings highlight that enhanced control measures could limit the future impact of HPAI on the poultry industry and reduce the risk of broader health threats.

## Introduction

Highly pathogenic avian influenza (HPAI) poses a significant ongoing threat to the poultry industry. Since 2020, the emergence of a reassorted genotype of H5N1 viruses within clade 2.3.4.4b has been associated with widespread outbreaks of HPAI in both wild birds and poultry worldwide [1]. The currently circulating H5N1 clade 2.3.4.4b strain has a significant fitness advantage over previously circulating viruses [2] and has impacted a larger number of bird species, notably causing large mortality events in seabird colonies [3, 4]. Moreover, spillover into mammals, including evidence of mammal-to-mammal transmission, such as in cattle in the USA [5, 6], as well as an increasing number of confirmed human infections [7, 8] indicate the zoonotic potential of the virus and the potential risk of a future pandemic occurring [9].

In Great Britain, there have been annual epizootic events affecting poultry premises since 2020, with hundreds of infected premises (IPs) during the 2022–23 season (1 October 2022 –– 30 September 2023) [10]. This has resulted in the culling of millions of birds to prevent further spread of infection, at substantial economic cost [2, 11]. The outbreaks traditionally followed a seasonal pattern, driven by the arrival of infection from migratory wild birds in the autumn and winter months, with fewer infections in poultry during the summer [12]. However, with a broader range of wild bird host species infected and endemic circulation in these resident wild birds [13], there has been an increased persistence in observed IPs over the summer months [10].

Most reported IPs were likely associated with spillover from local wild bird populations in 2022–23 [14], while infected migratory birds commonly increase the geographical spread of infection [15]. The majority of transmission to poultry has generally been attributed to wild ducks, geese, and gulls and in most cases is due to environmental contamination of infected faecal matter in water sources or direct contact with infected carcasses [16]. In the poultry premises, chickens and turkeys are typically highly susceptible to HPAI compared to duck species [17], although some studies suggest that circulating H5N1 clade 2.3.4.4b may be particularly well adapted to ducks [18, 19].

Premises-to-premises transmission has been reported in Europe [20], but there is little evidence that this is common in Great Britain. Phylogenetic analyses have identified premises-to-premises transmission as being likely for only a few select IPs [21]. Where premises-to-premises transmission does occur, it is likely due to the movement of vehicles or shared equipment between premises [22, 23]. Airborne transmission between premises is unlikely since evidence suggests that airborne particles containing HPAI virus can only travel very short distances (up to 10 metres) [24]. With that understanding of the likely modes of transmission, biosecurity measures are therefore essential to prevent the movement of HPAI virus onto premises. Biosecurity measures can include increased disinfection and cleaning, management and treatment of water, the prevention of wild-bird access to housing and feed storage, changing of footwear for poultry workers moving between premises, and improved fencing [25, 26]. In Great Britain, Avian Influenza Prevention Zones (AIPZ) have been introduced that legally require poultry owners to follow strict biosecurity measures and can include mandatory housing orders [27, 28]. As of March 2025, the use of HPAI vaccines for poultry is not permitted in Great Britain [29].

In the 2022–23 season, there was a national AIPZ in place from 17 October 2022 to 4 July 2023 [27], with a regional AIPZ in the counties of Norfolk, Suffolk, and parts of Essex before this from 27 September 2022, and so covering almost all the infections on premises seen during the season. A national housing order was also in place from 7 November 2022 to 18 April 2023 [28], with this beginning regionally in the East of England from 12 October 2022. Additional 3 km Protection Zones were established around IPs to limit transmission, alongside 10 km (7 km in addition to the 3 km) Surveillance Zones with increased record keeping and monitoring for infections [30]. For our mathematical analysis, we choose to model this season.

Mathematical models for HPAI outbreaks amongst poultry premises have been used to estimate the probability that large outbreaks occur in a variety of geographical settings and to identify areas at high risk of infection. In the context of Great Britain, spatial models have historically helped to determine the probability of outbreak clusters [31, 32]. Spatial modelling studies have also identified key parameter values for model simulations in other settings, such as Bangladesh [33] and Thailand [34]. Modelling approaches have been developed to produce risk maps of HPAI H5N1 Clade 2.3.4.4b spillover in Europe [35] and the USA [36]. Other studies have sought to identify suitable control policies for HPAI including vaccination, ring culling, increased surveillance, and contact tracing in countries such as Vietnam [37], Bangladesh [38], South Korea [39], and France and the Netherlands [40]. However, no known studies have been used to infer the transmission dynamics of recent outbreaks of HPAI in Great Britain, or to assess the impact of potential control strategies.

In this manuscript, we extended the work of previous modelling approaches, such as Jewell et al. [31] and Hill et al. [33], to adapt a spatial individual poultry premises-based model for Great Britain. We used this model to capture the infection dynamics in poultry premises across the 2022–23 season, the season with the largest number of HPAI-infected poultry premises (at the time of writing in March 2025). We used Markov chain Monte Carlo (MCMC) to parameterise the model using notification data from the 2022–2023 season, evaluating the quality of the model fit with model simulations and fitting statistics. We then consider the impact of biosecurity measures or the potential use of vaccines by implementing enhanced control measures in the immediate area surrounding an IP. Using model simulations, we explore how varying the stringency, duration, and area of the enhanced control zone affects the number of reported IPs. These results identify plausible mechanisms for HPAI transmission in poultry within a season in Great Britain and quantify the impact of additional control measures for a range of possible scenarios.

## Methods

### Data

We obtained demographic data on poultry premises in Great Britain and poultry case data from the Animal and Plant Health Agency. The demographic data included the centroid of each premises polygon (defined as a CPH — County/Parish/Holding number — entity), as well as the number of poultry reported as kept by species. We had demographic data from 1 December 2022, which falls within our fitting period. To provide the required inputs into the spatial model, we aggregated these flock counts into three categories for each premises: Galliformes (chickens, turkeys, etc.), waterfowl (ducks, geese, etc.), and other birds. We chose these categories to limit the number of bird types considered for a reasonable number of parameters in the model while allowing for differences in the transmission and susceptibility characteristics of different species. There were 48660 premises within our data set. While we assumed this was a complete list of all poultry in the country, some premises of any size may be missing from our data set and, in particular, we noted that it was not a legal requirement for premises with fewer than 50 captive birds to officially register their birds in the considered time period (before 1 October 2024), indicating that small premises may be missing [41].

In this study, we considered premises with H5N1-infected birds within the 2022–23 epidemiological season: 1 October 2022 – 30 September 2023. This period contains a peak in infections in the autumn/winter, with relatively fewer cases in the summer months, consistent with the known seasonality in infections [12]. There were 200 premises with confirmed H5N1 infection in the 2022–23 season (Figure 1).

**Figure 1:**
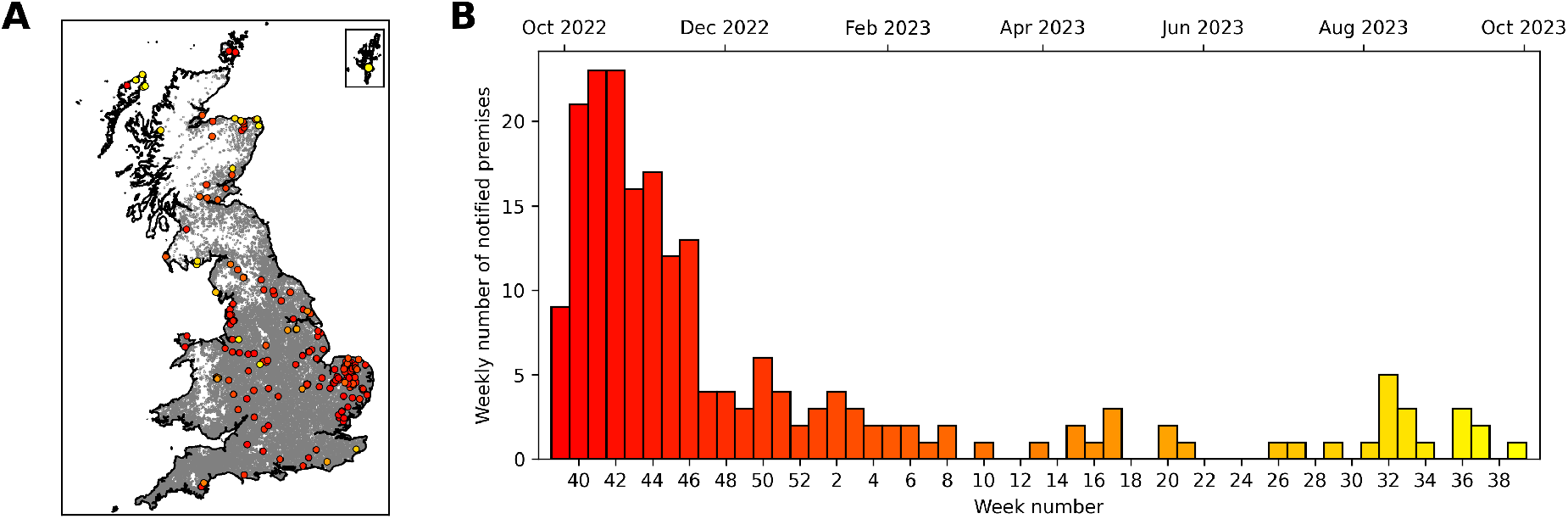
Reported cases of H5N1 in poultry premises in Great Britain for the 2022–23 season, 1 October 2022 – 30 September 2023. (A) A map of all poultry premises in the poultry keeper register on 1 December 2022 in Great Britain (grey dots), and the locations of infected premises (IPs) within this time period (coloured dots). The colour of the IP dots indicates the time within the season that the notification occurred, with red dots at the start of the season and yellow dots at the end. The inset map shows the Shetland Islands to the north of mainland Scotland. Source of map boundaries: Office for National Statistics licensed under the Open Government Licence v.3.0. The shapefiles used can be found at https://geoportal.statistics.gov.uk/datasets/5a393192a58a4e50baf87eb4d64ca828_0/explore. (B) A bar chart of the number of notified poultry premises in each week of the 2022–23 season. The colour scale of the bars indicates the progression of time, as shown in (A). The secondary axis on the top shows the start of selected months in this time period.

At any point in time, we assume that all IPs are included in the data set (due to identification by poultry exhibiting clinical signs or suffering mortality), unless they have yet to be reported. The IPs will be reported because HPAI is a notifiable disease in the UK, and poultry owners are legally obliged to report suspected infections to the Animal and Plant Health Agencies (APHA) [42]. Upon reporting or ‘notification’, a veterinary inspector typically visits the premises to test for the presence of HPAI within the poultry, where on disease confirmation, all susceptible poultry will be culled unless specific exemption criteria apply [43]. Additional measures such as movement bans, disinfection, and other restrictions may also be put in place.

We link data on IPs to the poultry premises register data by matching the coordinates of the premises with reported infections to the closest premises with similar reported poultry numbers. Full details are provided in the supplementary material. This means we have data on all premises contained in the poultry keeper register data for Great Britain on 1 December 2022, and for those premises where H5N1 infection was detected, the dates that this occurred before all birds were culled on the premises.

### Model

Our model is formulated as a discrete-time, individual-based, spatially explicit compartmental model for individual poultry premises. Our model conceptualisation uses a similar structure to models presented in previous studies on HPAI and other livestock diseases [33, 44–46]. At a given time, each premises can be in one of five given states: susceptible to infection S, exposed to infection E, infected and able to transmit infection I, notified as infected N, and removed by culling R. For each premises *i*, we denote *E*_*i*_ as the time at which the premises became exposed. This similarly applies for time of infection (*I*_*i*_), time of notification (*N*_*i*_), and time of culling (*R*_*i*_). We assume the infection events occurred according given infection rate *λ* _*j*_ (*t*).

The infectious pressure on premises *j* is given by:

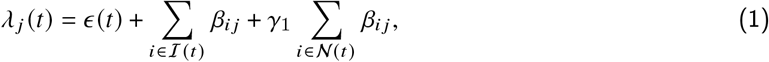

where *ℐ* (*t*) and *𝒩* (*t*) are the sets of premises infected and notified at time *t* respectively. This describes two components of infection: (i) a time-varying term, *ϵ* (*t*), for the background infection directly caused by spillover from wild bird populations; (ii) local infections in regions close to other poultry premises that are currently infectious (infected I or notified N) [47], captured by the two summation terms in Equation 1. The local force of infection from premises *i* to premises *j* is given by *β*_*ij*_ and *γ*_1_ is a scaling of the force of infection for premises that have been notified as infectious, as opposed to infectious but not yet notified. The local infection component from other poultry premises could be directly from other poultry premises, indirectly by intermediate wild bird infection or infectious virus on shared sites or equipment, or infection in other premises in close proximity could be indicative of an increased presence of H5N1 in local wild bird populations. Therefore, both components may be due to wild bird spillover, with known poultry infections spatially indicating potential higher-risk areas.

The constituent terms of *λ* _*j*_ (*t*) are further broken down to describe their functionality within the model. The background infection term:

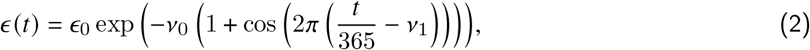

shows seasonal variation in its influence peaking on day *ν*_1_ in each year [12]. The force of infection on premises *j* by premises *i* is given by:

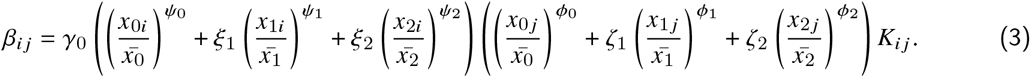

The force of infection *β*_*ij*_ therefore consists of four components. A multiplicative factor *γ*_0_; the infectivity of the premises *i*, which is the sum of the number of birds of the three species types (*x*_*ki*_, *k* = 0, 1, 2 for the three species types) divided by the mean value of the number of birds of each type across all premises 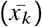 to the infectivity exponent (*ψ*_*k*_) and multiplied by infectivity factor (*ξ*_*k*_, where *ξ*_0_ = 1); the susceptibility of premises *j*, which takes a similar form to the infectivity, but with susceptibility exponents (*ϕ*_*k*_) and factors(*ζ*_*k*_); and the transmission kernel *K*_*ij*_, which is a function of distance between the infected and susceptible premises.

We include exponents on the number of birds in the infectivity and susceptibility components because previous studies have found that non-linear terms provide a better fit to epidemic data [48]. In this study, we also assume that the kernel takes a Cauchy form with:

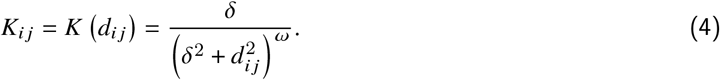

Therefore, the force of infection *λ* _*j*_ (*t*) is only spatially dependent on the distance to other infected and notified premises, where there is an additional multiplicative scaling parameter *γ*_1_ applied to the force of infection from notified premises. We also consider an exponential kernel in the supplementary material (SI).

Given a timestep [*t, t* + *δt*) where *δt* = 1 day, the probability that a susceptible premises becomes exposed on a given day is given by:

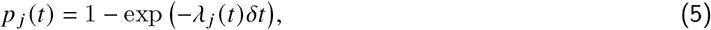

and from this exposure event, we then assume the infection progresses in discrete time through the infection states after a specified number of days. We fix the latent period and time to culling from notification for improved model tractability, but leave the time to notification as variable for each premises in the fitting process, since we expect this will have the most variance in reality due to differences in premises-level surveillance and visibility of symptoms in birds.

In further detail, we fix the time spent in the exposed class as four days before moving to the infected class, since the incubation period is generally less than seven days [49], but there is little specific literature on between-flock latency periods [50]. We then take the time in the infected class, before notification, as a random number of days given by *N* _*j*_ − *I* _*j*_ ∼ Gamma (4, 2). Gamma (*a, b*) describes the Gamma probability distribution with shape *a* and scale *b*. This allows for individual differences dependent on the specific premises, but provides an estimate that falls within the typical distribution [50]. The time from notification to culling the birds is taken as a fixed three days, which is consistent with the average report and confirmation dates in the data set.

### Model parameters and fitting

The model parameter values are determined using a Bayesian inference framework, where the parameters are updated using adaptive Metropolis–Hastings in a Markov Chain Monte Carlo method [51, 52]. This method has been implemented similarly in several studies [31, 45, 46, 53].

We fit sixteen model parameters with one fixed parameter *ω* = 1.3 to give the shape of our transmission kernel. More details are given in the supplementary information. See Table 1 for full details on all the model parameters. Prior distributions for these parameters are chosen such as to provide variance around a mean value elicited from expert opinions, or to be uninformed.

**Table 1:** List of model parameters with descriptions. Prior distributions are given for fixed parameters where Gamma (*a, b*) is a gamma distribution with shape *a* and scale *b* and Beta (*a, b*) is a beta distribution with shape parameters *a* and *b*. Note that *ξ*_0_ = 1 and *ζ*_0_ = 1 are not listed as *ξ*_*k*_ and *ζ*_*k*_ are relative to these parameters respectively for *k* = 1, 2.

| Parameters | Description | Prior/Value |
| --- | --- | --- |
| $\epsilon_0$ | Baseline infectious pressure | Gamma(1, $1 \times 10^{-5}$) |
| $\gamma_0$ | Infectious pressure contribution from IPs | Gamma(1, 0.01) |
| $\gamma_1$ | Multiplicative factor for infectious pressure contribution for notified premises | Gamma(1, 0.8) |
| $\delta$ | Decay of transmission rate between premises in the transmission kernel | Gamma(2, 1) |
| $[\psi_0, \psi_1, \psi_2]$ | Exponential terms in infectious pressure for infectious premises | Beta(2, 2) |
| $[\phi_0, \phi_1, \phi_2]$ | Exponential terms in infectious pressure for susceptible premises | Beta(2, 2) |
| $[\xi_1, \xi_2]$ | Relative transmissibility of species types | Gamma(1, 1) |
| $[\zeta_1, \zeta_2]$ | Relative susceptibility of species types | Gamma(1, 1) |
| $\nu_0$ | Shape of seasonality | Gamma(2, 2) |
| $\nu_1$ | Timing of seasonality | Beta(2, 2) |
| $\omega$ | Exponent in transmission kernel | 1.3 |
| $a$ | Shape of gamma distribution for time to notification | 4 |
| $b$ | Scale of gamma distribution for time to notification | 2 |

Also included within the fitting algorithm is a reversible-jump update, which allows the addition (and removal) of undetected or ‘occult’ infections within the model to assess how their presence changes the likelihood [45]. In this step, which occurs after each update to the parameter values, a fixed number of additional model alterations are considered: the change of the time to notification for a given premises, the addition of a new IP as an ‘occult’ infection that is yet to be notified (and so appear in the data set), and the removal of any previously added ‘occult’ infections.

### Model projections and enhanced control strategies

The fitted model can be used to stochastically simulate the epidemic from a given set of initially infected premises to compare back to the observed data. For the 2022–23 season, our initial conditions are a fixed set of 31 infected (exposed, infected, or notified) premises in all simulations across East of England (23 IPs), West Midlands (2), North West (2), East Midlands (1), South West (1), South East (1) and Scotland (1). The numbers were determined from our data set. The simulations are performed using a tau-leaping algorithm [54] in a grid-based system for computational speed [55, 56].

Additionally, we perform counterfactual simulations based on alternative scenarios using the same methodology. These model simulations specifically investigate how enhanced control might be enacted in response to an IP. We assume a baseline level of biosecurity, which is determined by the fitting process for the 2022–23 season, where in reality an AIPZ and housing order were in place for the majority of the season. Enhanced control measures could include increased cleaning and disinfection, and reduced risk of contact with wild birds and contamination of water sources, feed storage and housing, and the potential use of vaccination. All these measures will have the effect in the model of reducing the susceptibility of the poultry that could become infected with HPAI, and so the risk of HPAI incursion.

In our model simulation strategies, enhanced control will be mandated on the discovery of the presence of HPAI infection within a premises from the time of culling and will last for a specified number of days. This will either occur in all poultry premises within a particular radius (5 km, 10 km, or 15 km) of the IP or within all poultry premises in the same county or region (see supplementary information for full details on Great British counties and regions). The effect of enhanced control is to reduce the susceptibility to infection of the nearby premises due to the reduced risk from the improved control measures. In this manuscript, we consider enhanced control measures that reduce susceptibility to 80%, 60%, 40%, and 20% of what the susceptibility was determined to be in model fitting (here referred to as a susceptibility factor of 0.8, 0.6, 0.4, or 0.2) for 7, 14, 21 or 28 days since the date of culling on an IP.

## Results

### Verifying the model fitting

The MCMC process was successful in providing posterior parameter distributions for each fitted parameter, with good convergence of the chains (see supplementary material for full details).

Sampling from the joint posterior distributions to simulate the model forward in time from the observed initial conditions on 1 October 2022, we verify that we achieve a good correspondence back to the data (Figure 2). The data points for the weekly number of premises reported as infected fall within the 95% prediction intervals of the model simulations, with the closest matching model simulations providing a strong temporal fit to the data (Figure 2A).

**Figure 2:**
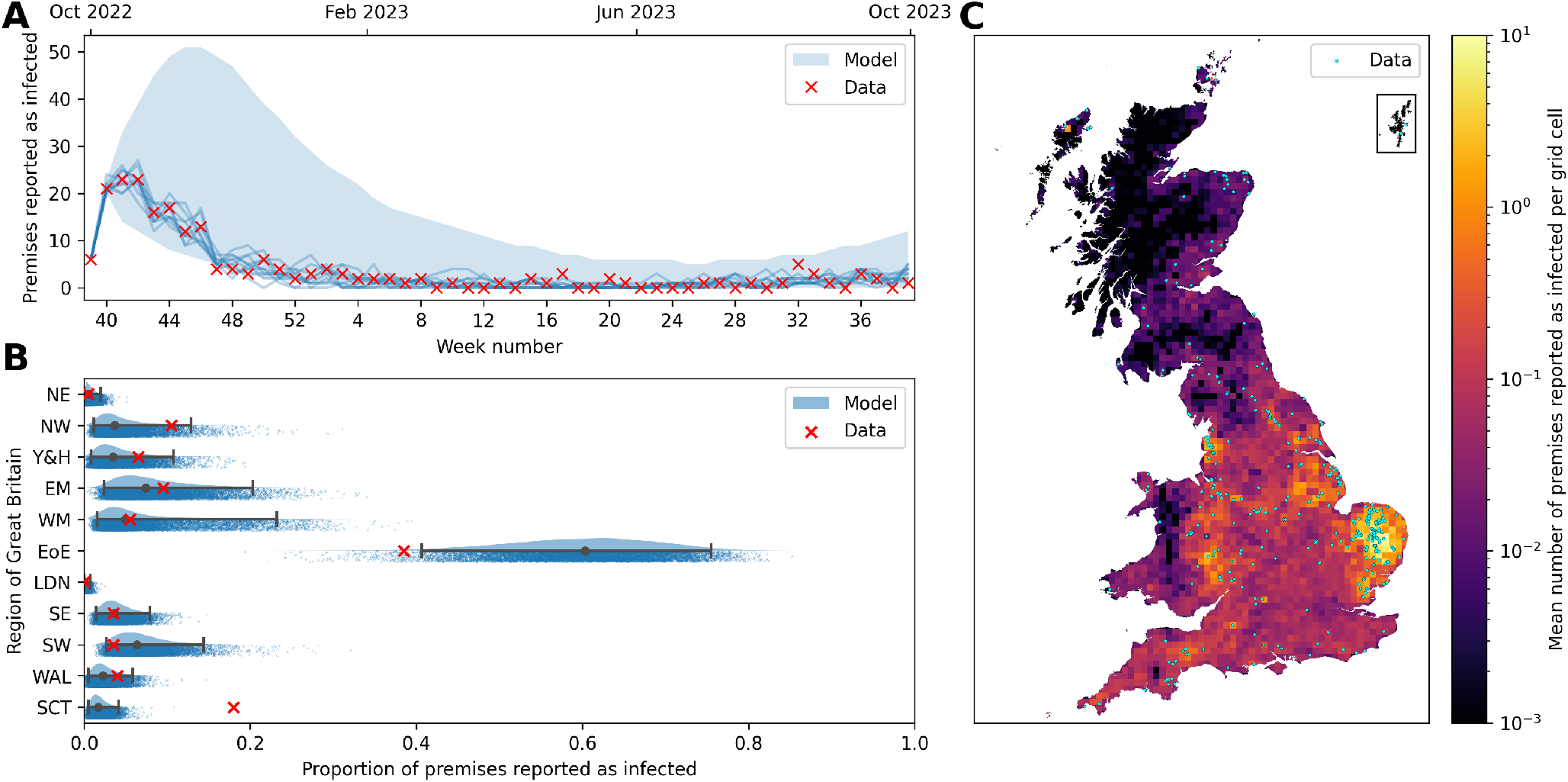
Comparison of model simulations to premises notification of infection data for 1 October 2022 – 30 September 2023. (A) Time series of the weekly number of premises with reported infections. The red crosses indicate the data, while the shaded blue regions show the 95% prediction intervals of stochastic model simulations. Blue lines indicate the best fitting 10 individual realisations of the simulation out of the total 10,000. (B) Raincloud plots [57] for the proportion of infected premises (IPs) across the eleven geographical regions of Great Britain. The data are represented by red crosses, while the central prediction interval and rainclouds show model simulations. The grey dot shows the median value, and the whiskers give the full 95% prediction interval. Above each interval is a half-violin plot of the distribution of the simulations, and below is a jittered scatter of each individual simulation. The names of the regions in full are: NE – North East, NW – North West, Y&H – Yorkshire and the Humber, EM – East Midlands, WM – West Midlands, EoE – East of England, LDN – London, SE – South East, SW – South West, WAL – Wales, SCT – Scotland. (C) Map of Great Britain divided into 10 km × 10 km grid cells coloured to show the mean number of IPs in each cell for the 2022–23 season. Blue dots overlaying the grid cells show the locations of the true IPs in this season. Source of map boundaries: Office for National Statistics licensed under the Open Government Licence v.3.0. The shapefiles used can be found at https://geoportal.statistics.gov.uk/datasets/5a393192a58a4e50baf87eb4d64ca828_0/explore.

To consider the spatial model fit, we divide the number of IPs from the national model simulations into the eleven geographical regions of Great Britain. We observe that we have achieved a favourable match to the data spatially in most of these regions, with the proportion of the total IPs within the regions having a close median value in simulations to the real-world data for the season (Figure 2B). The two notable exceptions to this occur in the East of England and Scotland. For the East of England, we predict a larger proportion of cases than observed, with the inverse true of Scotland. This can, in part, be explained by the initial conditions of the model simulations, as there happen to be initially many IPs in the East of England and few in Scotland, as per the data.

Therefore, we highlight the difficulty of achieving an optimal model fit to this data set, both spatially and temporally simultaneously, due to the large state space of possible outcomes and relying on random spillover events in poultry premises to recreate the observed localised outbreaks seen in the data set. This is despite demonstrating that the model can generate simulations with comparable results to the original data. This overall quality of the model fit is shown on the map in Figure 2C. The blue dots, which represent the IPs in the data set, typically fall within the 10 km × 10 km grid cells that are predicted to be at the highest risk of infection in the model.

We additionally calculate that there will likely be few ‘occult’ infections with a median number of 2 IPs identified in model fitting, with the true value in the data of 0 falling within the prediction intervals. Further details on the ‘occult’ infections and the quality of the model fit are presented in the supplementary material.

### Control strategy scenarios

Model projections for the effect of enhanced control on the total number of IPs show that improvements may be possible by a concerted effort to reduce the potential for transmission in the vicinity of premises that have previously detected infection. If biosecurity can be improved or vaccines delivered, such that premises within an enhanced control zone that lasts for 21 days are 40% less likely to be infected (susceptibility factor = 0.6), then the median number of IPs nationally can be reduced by up to 53% (depending on the size of the enhanced control zone) (Figure 3). However, a susceptibility factor of 0.8 can still reduce the median number of IPs by up to 35% as well as a reduction in the uncertainty.

**Figure 3:**
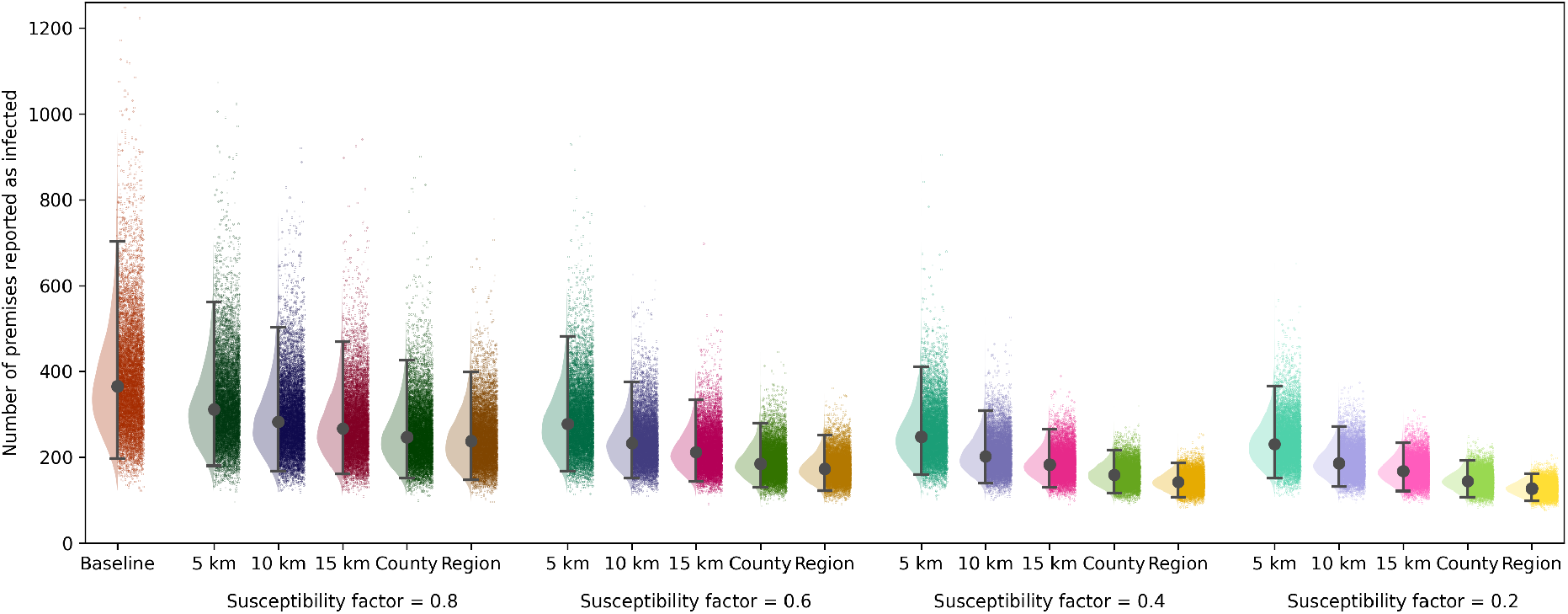
Projected impact of enhanced control. Each raincloud plot [57] shows the total number of infected premises (IPs) in the season for a particular enhanced control scenario, based on the region in which premises are affected (with a 5 km, 10 km, or 15 km radius, or across the county or the region) and the reduction in susceptibility due to improved control, which is here called the susceptibility factor. Estimates are obtained from 10,000 model simulations for each scenario. A susceptibility factor of 1 indicates no change in susceptibility, whereas the numbers less than 1, show multiplicative factors that result in reduced susceptibility to HPAI infection. In each raincloud, it is assumed that enhanced control measures affect the given region for 21 days after poultry has been culled on the IPs and premises within multiple impacted regions have reduced susceptibility according only to the most recent IP. Within the rainclouds, we show the median simulation value (grey dots), and 95% prediction interval (whiskers), alongside a half violin plot of the distribution (to the left) and jittered scatter of all simulations (to the right).

The size of the enhanced control zone has a large impact on the efficacy of the scenario, with the additional 5 km zones only causing a modest reduction in the median number of IPs (27% reduction), even when the susceptibility factor is 0.2. Increasing this radius to 10 km or 15 km impacts a much larger number of premises and so results in a larger reduction in the number of IPs, particularly when combined with lower susceptibility factors. Scaling the zone up to the full county or region shows further improvements in reducing the number of IPs. This indicates it is likely insufficient to solely consider the immediate area around an IP, likely due to the movement of wild birds. Larger zones for either strict to moderate control, with increased surveillance, could improve the reduction in infection. In particular, greatly enhanced control (susceptibility factor = 0.2) combined with enacting this over the full geographical region both greatly reduce the median number of IPs as well as the uncertainty in this number, eliminating some of the worst-case scenarios.

The duration of enhanced control has a relatively smaller impact on the number of IPs across the season than the susceptibility factor or size of the zone (Figure 4). There is very little difference in IPs for the season when the duration of enhanced control is varied between 7 and 28 days. This emphasises that most of the impact is due to quickly implementing the enhanced control zone upon detection of infection, with diminishing returns for keeping the zone in place for many weeks. Secondary premises will most likely become infectious close to when the original IP is detected. The largest impact of the duration occurs when the susceptibility factor is small (such that the enhanced control is having a large effect) and the zone area is also small. In this scenario, there is a large benefit in terms of susceptibility for local premises being within the zone, but local, as yet undetected IPs may fall outside this small area, and so increasing the duration of the enhanced control zone prevents more infections.

**Figure 4:**
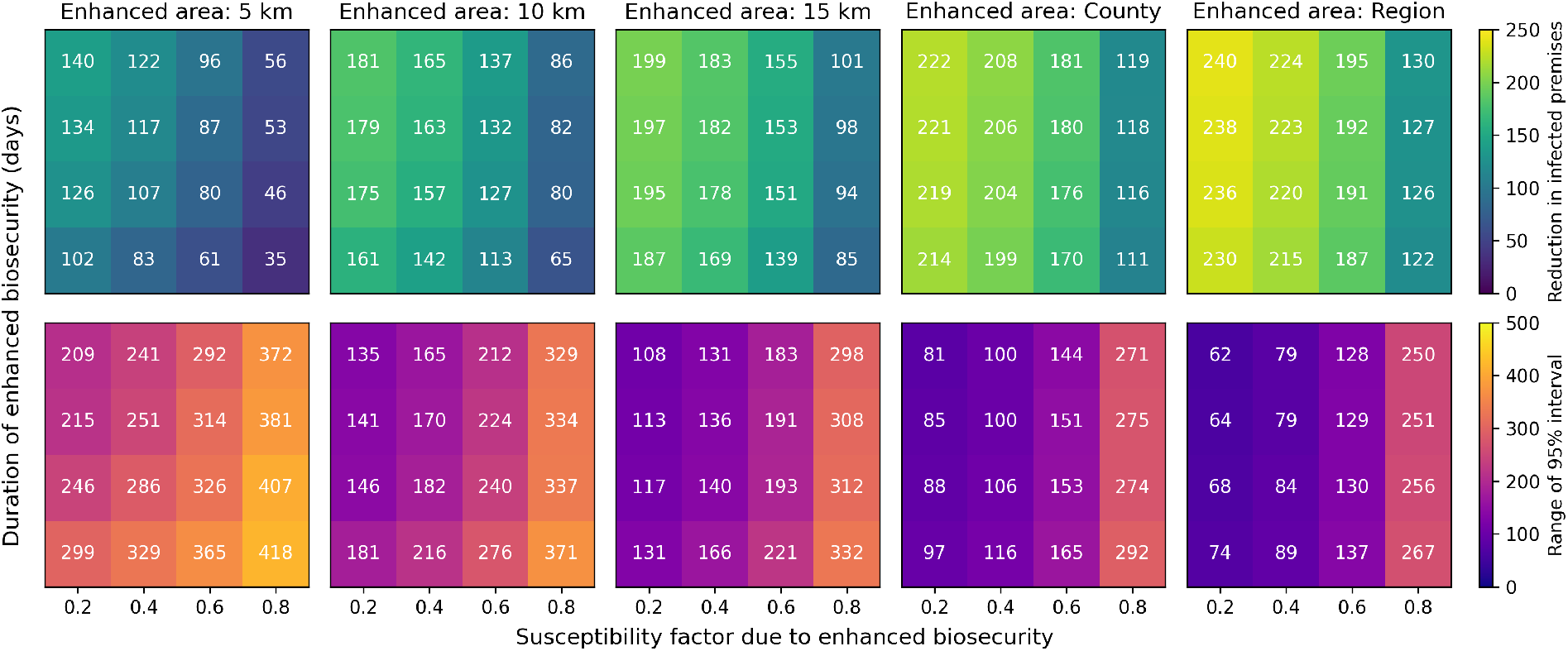
Heat maps of the median reduction in number of infected premises (IPs) and 95% prediction interval range under different scenarios. The top row of panels shows the median reduction in IPs for a given enhanced control duration, susceptibility factor, and zone. The bottom row shows the respective 95% prediction interval range. Estimates are obtained from 10,000 model simulations for each scenario. Each panel shows the results for different enhanced control zone areas.

The largest uncertainties occur when the susceptibility factor is large because then there is little impact of the additional control measures, and this scenario is closest to no control. Similarly, there is also more uncertainty when the control zone is smaller. Stringent control measures, coupled with implementing them across larger areas, limit the potential for large outbreaks to occur.

## Discussion

Overall, this study showcases a spatial model for HPAI infection in the poultry of Great Britain that has been newly adapted from previous livestock modelling studies [33, 44–46] for use with H5N1 clade 2.3.4.4b. The model is both shown to capture the observed IP data for the 2022–23 season, but could also be applied to other time periods to similarly estimate the underlying infection dynamics. The complex fitting process is computationally intensive; given our data on IPs the inference procedure identifies the most probable distributions for our model parameters. We also used the model to determine the most likely as yet undetected IPs by the end of our data period (or the ‘occult’ infections), such that the updated transmission process by the inclusion of these infections most improves our model fit.

We have emphasised that our mechanistic modelling approach can be additionally used to perform counter-factual simulations for the season. We chose to consider enhanced control measures to limit the potential for further transmission events within a certain area. These simulations indicate that additional measures around premises that have had confirmed infections can greatly reduce the total number of premises that experience infections. The greatest impact occurs when enhanced control is established in the premises within a large area centred on an IP. There is a substantial reduction of additional IPs, even if just the immediate premises within a 5 km radius, make efforts to reduce the potential for HPAI transmission. We note that if spillover from wild birds is the primary route by which poultry become infected, biosecurity measures to minimise contact with wild birds are of critical importance.

Our results therefore align with the national policy that there is a benefit to implementing increased biosecurity in a radius around IPs [30]. However, we have highlighted there is an additional benefit to increasing the size of this radius (Figure 3), although clearly this will come at an additional cost. Vaccinations could also offer a solution to reduce the susceptibility of poultry and hence reduce the number of IPs, although there are practical and commercial issues with delivering vaccinations, as well as increased costs [29]. In particular, the use of vaccinations would result in reduced mortality and fewer symptomatic HPAI infections in poultry, decreasing the probability of detection and increasing costs associated with surveillance [58].

We also note that there will be variations in how strictly control measures are applied, particularly in small-scale poultry premises. National surveys in the UK have indicated there are differences in the actions of small-scale poultry keepers in regards to both awareness and compliance with restrictions, as well as trust in authorities [11, 59]. Therefore, practically, it may be difficult to uniformly implement enhanced control strictly across all the required areas.

In designing our model, we have made the simplifying assumption that background infectious pressure from wild bird spillover into poultry premises is spatially uniform across Great Britain. This could be challenged by incorporating spatial information on wild bird habitats and detected cases into the *ϵ* term in the model equations (Equation 2). Alternatively, known environmental sources of infection or reported wild bird cases could be added as pseudo-premises to the model to include additional transmission sources. However, we have shown that we are able to achieve a remarkably good match to the real-world data for the 2022–23 season, given the lack of this information in the model (Figure 2).

In considering components of our spatial model fit, the biggest weakness is in matching the total number of IPs in the East of England in Scotland. However, this is related to the complexity of forward simulating a full year of HPAI transmission from the real-world initial conditions of 1 October 2022, alongside previously mentioned considerations. In the East of England, there were initially 23 IPs at the start of the season, and so a majority of model projections will favour continued transmission here, since there are many initially infected premises. Transmission within this region does not rely on new random introductions from wild bird spillover. Whereas in Scotland, the model starts with a single IP (on the relatively remote Isle of Lewis) and so relies on further chance model events of spillover in this region to generate the substantial number of infections observed across the region in the data. The discrepancy is therefore not a failure of model fitting (given the spatially invariant model parameter *ϵ*), but aims to match stochastic outputs across a long time period.

We have also assumed that the latent period and time to culling are fixed and the same for all IPs, in order to reduce the complexity of the fitting process. However, our chosen values are typical of the literature [50]. We have also chosen to omit continued transmission of infection in the area around an IP after culling the poultry to reduce our model complexity. Therefore, we may underestimate the benefit of continued enhanced control measures for longer periods after notification.

To further refine the model, we would like to understand the mechanisms behind transmission by untangling which transmission events come directly from wild bird spillover events versus those that originate from other infected poultry premises. This would enable us to better identify the risks associated with premises reporting infections. At present, the transmission kernel in our model includes many transmission routes: direct transmission between premises, transmission through an intermediary wild bird or birds, transmission through shared contaminated resources of poultry premises including poultry workers, and transmission directly from wild birds due to the area being a hotspot of HPAI infection, observed due to other local IPs.

Therefore, future adaptations of the model could involve considering industry links between poultry premises, using wild bird infection data, or considering phylogenetic analyses, albeit outside of the scope of the present study. In addition, if the current HPAI vaccination policy were to change to allow the vaccination of poultry, we could adapt our enhanced control strategy parameters to specifically consider vaccination, rather than only considering it as part of a package of interventions that could be used to reduce HPAI susceptibility in poultry [29].

Highly pathogenic avian influenza continues to be a worldwide concern due to the devastating impact on both poultry and wild bird populations, particularly in recent years, combined with the fact that further reassortment events could lead to the emergence of a virus with the potential to cause pandemics. For the recently circulating H5N1 virus in birds of Great Britain, we have demonstrated the capacity for outbreaks within the country and highlighted the potential for enhanced control measures to reduce the impact of HPAI outbreaks in the Great British poultry industry in the future.

## Data availability

Data on HPAI cases in poultry and Great British poultry premises demography are available on request from the Animal and Plant Health Agency (APHA).

## Code availability

The model code and scripts needed to generate the main figures can be found at https://github.com/cnd27/HPAI_control_measures/.

## Acknowledgements

The authors would like to thank Alexander Mastin and Ruth Moir from the Animal and Plant Health Agency (APHA) and Helen Roberts from the Department for Environment, Food and Rural Affairs (DEFRA) for the provision of demographic and case data used in this work. The work was supported by a joint EEID NSF/BBSRC grant [BB/X005224/1] and a BBSRC grant [BB/X016137/1].

## Author contributions

CND: Data curation, Formal analysis, Software, Validation, Visualisation, Writing-Original draft, Writing-Review and Editing. EMH: Methodology, Writing-Review and Editing. CPJ: Conceptualisation, Methodology, Writing-Review and Editing. KR: Writing-Review and Editing. RNT: Conceptualisation, Methodology, Writing-Review and Editing. MJT: Conceptualisation, Data curation, Methodology, Validation, Writing-Review and Editing.

## References

[1] Jing Yang, Chunge Zhang, Yue Yuan, Ju Sun, Lu Lu, Honglei Sun, Heting Sun, Dong Chu, Siyuan Qin, Jianjun Chen, et al. Novel avian influenza virus (H5N1) clade 2.3.4.4b reassortants in migratory birds, china. Emerging Infectious Diseases, 29(6):1244, 2023.

[2] Michelle Wille and Ian G Barr. Resurgence of avian influenza virus. Science, 376(6592):459–460, 2022.

[3] Johanna A Harvey, Jennifer M Mullinax, Michael C Runge, and Diann J Prosser. The changing dynamics of highly pathogenic avian influenza H5N1: Next steps for management & science in north america. Biological Conservation, 282:110041, 2023.

[4] Marco Falchieri, Scott M Reid, Craig S Ross, Joe James, Alexander MP Byrne, Madalina Zamfir, Ian H Brown, Ashley C Banyard, Glen Tyler, Emma Philip, et al. Shift in HPAI infection dynamics causes significant losses in seabird populations across great britain. Veterinary Record, 191(7):294–296, 2022.

[5] Thomas Peacock, Louise Moncla, Gytis Dudas, David VanInsberghe, Ksenia Sukhova, James O Lloyd-Smith, Michael Worobey, Anice C Lowen, and Martha I Nelson. The global H5N1 influenza panzootic in mammals. Nature, pages 1–2, 2024.

[6] Gabriele Neumann and Yoshihiro Kawaoka. Highly pathogenic H5N1 avian influenza virus outbreak in cattle: the knowns and unknowns. Nature Reviews Microbiology, 22(9):525–526, 2024.

[7] Isabel Oliver, Jonathan Roberts, Colin S Brown, Alexander MP Byrne, Dominic Mellon, Rowena DE Hansen, Ashley C Banyard, Joe James, Matthew Donati, Robert Porter, et al. A case of avian influenza a (H5N1) in england, january 2022. Eurosurveillance, 27(5):2200061, 2022.

[8] Shikha Garg, Katie Reinhart, Alexia Couture, Krista Kniss, C Todd Davis, Marie K Kirby, Erin L Murray, Sophie Zhu, Vit Kraushaar, Debra A Wadford, et al. Highly pathogenic avian influenza a (H5N1) virus infections in humans. New England Journal of Medicine, 2024.

[9] The Lancet Infectious Diseases. What is the pandemic potential of avian influenza a (H5N1)? The Lancet. Infectious diseases, 24(5):437, 2024.

[10] Megan Arter-Hazzard, Geraldine Burns, Candida Adridge, Paul Gale, Lauren Perrin, and Joe Bowen. Updated outbreak assessment #46: High pathogenicity avian influenza (HPAI) in the uk and europe. https://assets.publishing.service.gov.uk/media/652d4c0d697260000dccf8d5/HPAI_Europe46_10_October_2023.pdf. Accessed: 2025-01-30.

[11] Emma McClaughlin, Sol Elliott, Sarah Jewitt, Matthew Smallman-Raynor, Stephen Dunham, Tamsin Parnell, Michael Clark, and Rachael Tarlinton. Uk flockdown: A survey of smallscale poultry keepers and their understanding of governmental guidance on highly pathogenic avian influenza (HPAI). Preventive Veterinary Medicine, 224:106117, 2024.

[12] Jacintha GB Van Dijk, Bethany J Hoye, Josanne H Verhagen, Bart A Nolet, Ron AM Fouchier, and Marcel Klaassen. Juveniles and migrants as drivers for seasonal epizootics of avian influenza virus. Journal of Animal Ecology, 83(1):266–275, 2014.

[13] Alice Fusaro, Bianca Zecchin, Edoardo Giussani, Elisa Palumbo, Montserrat Agüero-García, Claudia Bachofen, Ádám Bálint, Fereshteh Banihashem, Ashley C Banyard, Nancy Beerens, et al. High pathogenic avian influenza a (h5) viruses of clade 2.3.4.4b in europe—why trends of virus evolution are more difficult to predict. Virus evolution, 10(1):veae027, 2024.

[14] Department for Environment, Food & Rural Affairs and Animal and Plant Health Agency. Highly pathogenic avian influenza h5n1 outbreaks in great britain: October 2022 to september 2023. https://assets.publishing.service.gov.uk/media/66570246dc15efdddf1a84cd/bird-flu-epi-report-2022-2023.pdf. Accessed: 2025-01-30.

[15] Dae-sung Yoo, Sung-Il Kang, Yu-Na Lee, Eun-Kyoung Lee, Woo-yuel Kim, and Youn-Jeong Lee. Bridging the local persistence and long-range dispersal of highly pathogenic avian influenza virus (HPAIV): a case study of HPAIV-infected sedentary and migratory wildfowls inhabiting infected premises. Viruses, 14(1):116, 2022.

[16] Artem Blagodatski, Kseniya Trutneva, Olga Glazova, Olga Mityaeva, Liudmila Shevkova, Evgenii Kegeles, Nikita Onyanov, Kseniia Fede, Anna Maznina, Elena Khavina, et al. Avian influenza in wild birds and poultry: dissemination pathways, monitoring methods, and virus ecology. Pathogens, 10(5):630, 2021.

[17] Anita Puranik, Marek J Slomka, Caroline J Warren, Saumya S Thomas, Sahar Mahmood, Alexander MP Byrne, Andrew M Ramsay, Paul Skinner, Samantha Watson, Helen E Everett, et al. Transmission dynamics between infected waterfowl and terrestrial poultry: Differences between the transmission and tropism of H5N8 highly pathogenic avian influenza virus (clade 2.3.4.4a) among ducks, chickens and turkeys. Virology, 541:113–123, 2020.

[18] Joe James, Elizabeth Billington, Caroline J Warren, Dilhani De Sliva, Cecilia Di Genova, Maisie Airey, Stephanie M Meyer, Thomas Lewis, Jacob Peers-Dent, Saumya S Thomas, et al. Clade 2.3.4.4b H5N1 high pathogenicity avian influenza virus (HPAIV) from the 2021/22 epizootic is highly duck adapted and poorly adapted to chickens. Journal of General Virology, 104(5):001852, 2023.

[19] Giulia Graziosi, Caterina Lupini, Elena Catelli, and Silvia Carnaccini. Highly pathogenic avian influenza (HPAI) H5 clade 2.3.4.4b virus infection in birds and mammals. Animals, 14(9):1372, 2024.

[20] Claire Guinat, Cecilia Valenzuela Agüí, Timothy G Vaughan, Jérémie Scire, Anne Pohlmann, Christoph Staubach, Jacqueline King, Edyta Świętoń, Adam Dan, Lenka Černíková, et al. Disentangling the role of poultry farms and wild birds in the spread of highly pathogenic avian influenza virus in europe. Virus Evolution, 8(2):veac073, 2022.

[21] Alexander MP Byrne, Joe James, Benjamin C Mollett, Stephanie M Meyer, Thomas Lewis, Magdalena Czepiel, Amanda H Seekings, Sahar Mahmood, Saumya S Thomas, Craig S Ross, et al. Investigating the genetic diversity of H5 avian influenza viruses in the united kingdom from 2020–2022. Microbiology spectrum, 11(4):e04776–22, 2023.

[22] Jennifer E Dent, Rowland R Kao, Istvan Z Kiss, Kieran Hyder, and Mark Arnold. Contact structures in the poultry industry in great britain: exploring transmission routes for a potential avian influenza virus epidemic. BMC Veterinary Research, 4:1–14, 2008.

[23] Dae-Sung Yoo, Byung chul Chun, Younjung Kim, Kwang-Nyeong Lee, and Oun-Kyoung Moon. Dynamics of inter-farm transmission of highly pathogenic avian influenza h5n6 integrating vehicle movements and phylogenetic information. Scientific Reports, 11(1):24163, 2021.

[24] Joe James, Caroline J Warren, Dilhani De Silva, Thomas Lewis, Katherine Grace, Scott M Reid, Marco Falchieri, Ian H Brown, and Ashley C Banyard. The role of airborne particles in the epidemiology of clade 2.3.4.4b H5N1 high pathogenicity avian influenza virus in commercial poultry production units. Viruses, 15(4):1002, 2023.

[25] Kathryn Glass, Belinda Barnes, A Scott, J-A Toribio, Barbara Moloney, Mini Singh, and Marta Hernandez-Jover. Modelling the impact of biosecurity practices on the risk of high pathogenic avian influenza outbreaks in australian commercial chicken farms. Preventive Veterinary Medicine, 165:8–14, 2019.

[26] Dae-Sung Yoo, Kwang-Nyeong Lee, Byung-Chul Chun, Ho-Sung Lee, Hyuk Park, and Jong-Kwan Kim. Preventive effect of on-farm biosecurity practices against highly pathogenic avian influenza (hpai) h5n6 infection on commercial layer farms in the republic of korea during the 2016-17 epidemic: A case-control study. Preventive Veterinary Medicine, 199:105556, 2022.

[27] Department for Environment, Food & Rural Affairs and Animal and Plant Health Agency. Avian influenza: Prevention zone declared across great britain. https://www.gov.uk/government/news/avian-influenza-prevention-zone-declared-across-great-britain. Accessed: 2025-03-03.

[28] Department for Environment, Food & Rural Affairs and Animal and Plant Health Agency. Avian influenza: Housing order to be introduced across england. https://www.gov.uk/government/news/avian-influenza-housing-order-to-be-introduced-across-england. Accessed: 2025-03-03.

[29] Department for Environment, Food & Rural Affairs, Animal and Plant Health Agency, and Veterinary Medicines Directorate. Vaccination of birds against high pathogenicity avian influenza (bird flu) joint statement from the avian influenza vaccination taskforce. https://www.gov.uk/government/publications/vaccination-of-birds-against-high-pathogenicity-avian-influenza-bird-flu-joint-statement. Accessed: 2025-03-25.

[30] Department for Environment, Food & Rural Affairs and Animal and Plant Health Agency. Bird flu: rules in disease control zones in england. https://www.gov.uk/guidance/avian-influenza-bird-flu-cases-and-disease-control-zones-in-england. Accessed: 2025-03-03.

[31] Chris P Jewell, Theodore Kypraios, RM Christley, and Gareth O Roberts. A novel approach to real-time risk prediction for emerging infectious diseases: a case study in avian influenza H5N1. Preventive veterinary medicine, 91(1):19–28, 2009.

[32] Kieran J Sharkey, Roger G Bowers, Kenton L Morgan, Susan E Robinson, and Robert M Christley. Epidemiological consequences of an incursion of highly pathogenic H5N1 avian influenza into the british poultry flock. Proceedings of the Royal Society B: Biological Sciences, 275(1630):19–28, 2008.

[33] Edward M Hill, Thomas House, Madhur S Dhingra, Wantanee Kalpravidh, Subhash Morzaria, Muzaffar G Osmani, Mat Yamage, Xiangming Xiao, Marius Gilbert, and Michael J Tildesley. Modelling H5N1 in bangladesh across spatial scales: Model complexity and zoonotic transmission risk. Epidemics, 20:37–55, 2017.

[34] Renata Retkute, Chris P Jewell, Thomas P Van Boeckel, Geli Zhang, Xiangming Xiao, Weerapong Thanapongtharm, Matt Keeling, Marius Gilbert, and Michael J Tildesley. Dynamics of the 2004 avian influenza H5N1 outbreak in thailand: the role of duck farming, sequential model fitting and control. Preventive veterinary medicine, 159:171–181, 2018.

[35] Sarah Hayes, Joe Hilton, Joaquin Mould-Quevodo, Christl Donnelly, Matthew Baylis, and Liam Brierley. Ecology and environment predict spatially stratified risk of highly pathogenic avian influenza in wild birds across europe. bioRxiv, pages 2024–07, 2024.

[36] Diann J Prosser, Cody M Kent, Jeffery D Sullivan, Kelly A Patyk, Mary-Jane McCool, Mia Kim Torchetti, Kristina Lantz, and Jennifer M Mullinax. Using an adaptive modeling framework to identify avian influenza spillover risk at the wild-domestic interface. Scientific Reports, 14(1):14199, 2024.

[37] Patrick GT Walker, Simon Cauchemez, Raphaëlle Métras, Do Huu Dung, Dirk Pfeiffer, and Azra C Ghani. A bayesian approach to quantifying the effects of mass poultry vaccination upon the spatial and temporal dynamics of H5N1 in northern vietnam. PLoS computational biology, 6(2):e1000683, 2010.

[38] Edward M Hill, Thomas House, Madhur S Dhingra, Wantanee Kalpravidh, Subhash Morzaria, Muzaffar G Osmani, Eric Brum, Mat Yamage, Md A Kalam, Diann J Prosser, et al. The impact of surveillance and control on highly pathogenic avian influenza outbreaks in poultry in dhaka division, bangladesh. PLoS computational biology, 14(9):e1006439, 2018.

[39] Dae-Sung Yoo, Byungchul Chun, Kyung-Duk Min, Jun-Sik Lim, Oun-Kyoung Moon, and Kwang-Nyeong Lee. Elucidating the local transmission dynamics of highly pathogenic avian influenza h5n6 in the republic of korea by integrating phylogenetic information. Pathogens, 10(6):691, 2021.

[40] Sébastien Lambert, Lisa Fourtune, Peter HF Hobbelen, Julie Baca, José L Gonzales, Armin RW Elbers, and Timothée Vergne. Optimizing contact tracing for avian influenza in poultry flocks. Journal of the Royal Society Interface, 22(222):20240523, 2025.

[41] Department for Environment, Food & Rural Affairs, The Scottish Government, and Welsh Government. New measures to help protect poultry industry from bird flu. https://www.gov.uk/government/news/new-measures-to-help-protect-poultry-industry-from-bird-flu. Accessed: 2025-01-30.

[42] Department for Environment, Food & Rural Affairs and Animal and Plant Health Agency. Notifiable diseases in animals. https://www.gov.uk/government/collections/notifiable-diseases-in-animals. Accessed: 2025-01-30.

[43] Sarah Jewitt, Emma McClaughlin, Sol Elliott, Matthew Smallman-Raynor, Michael Clark, Stephen Dunham, and Rachael Tarlinton. Veterinarians’ knowledge and experience of avian influenza and perspectives on control measures in the uk. Veterinary Record, page e3713, 2024.

[44] Matt J Keeling, Mark EJ Woolhouse, Darren J Shaw, Louise Matthews, Margo Chase-Topping, Dan T Haydon, Stephen J Cornell, Jens Kappey, John Wilesmith, and Bryan T Grenfell. Dynamics of the 2001 uk foot and mouth epidemic: stochastic dispersal in a heterogeneous landscape. Science, 294(5543):813– 817, 2001.

[45] Chris P Jewell, Theodore Kypraios, Peter Neal, and Gareth O Roberts. Bayesian analysis for emerging infectious diseases. Bayesian Analysis, 4(3):465–496, 2009.

[46] William JM Probert, Chris P Jewell, Marleen Werkman, Christopher J Fonnesbeck, Yoshitaka Goto, Michael C Runge, Satoshi Sekiguchi, Katriona Shea, Matt J Keeling, Matthew J Ferrari, et al. Real-time decision-making during emergency disease outbreaks. PLoS computational biology, 14(7):e1006202, 2018.

[47] Rob Deardon, Stephen P Brooks, Bryan T Grenfell, Matthew J Keeling, Michael J Tildesley, Nicholas J Savill, Darren J Shaw, and Mark EJ Woolhouse. Inference for individual-level models of infectious diseases in large populations. Statistica Sinica, 20(1):239, 2010.

[48] Michael J Tildesley, Rob Deardon, Nicholas J Savill, Paul R Bessell, Stephen P Brooks, Mark EJ Woolhouse, Bryan T Grenfell, and Matt J Keeling. Accuracy of models for the 2001 foot-and-mouth epidemic. Proceedings of the Royal Society B: Biological Sciences, 275(1641):1459–1468, 2008.

[49] Writing Committee of the Second World Health Organization Consultation on Clinical Aspects of Human Infection with Avian Influenza A (H5N1) Virus. Update on avian influenza a (h5n1) virus infection in humans. New England Journal of Medicine, 358(3):261–273, 2008.

[50] Carsten Kirkeby and Michael P Ward. A review of estimated transmission parameters for the spread of avian influenza viruses. Transboundary and Emerging Diseases, 69(6):3238–3246, 2022.

[51] Heikki Haario, Eero Saksman, and Johanna Tamminen. An adaptive Metropolis algorithm. Bernoulli, 7(2):223 – 242, 2001.

[52] Simon EF Spencer. Accelerating adaptation in the adaptive metropolis–hastings random walk algorithm. Australian & New Zealand Journal of Statistics, 63(3):468–484, 2021.

[53] Lloyd AC Chapman, Chris P Jewell, Simon EF Spencer, Lorenzo Pellis, Samik Datta, Rajib Chowdhury, Caryn Bern, Graham F Medley, and T Deirdre Hollingsworth. The role of case proximity in transmission of visceral leishmaniasis in a highly endemic village in bangladesh. PLoS neglected tropical diseases, 12(10):e0006453, 2018.

[54] Daniel T Gillespie. Approximate accelerated stochastic simulation of chemically reacting systems. The Journal of chemical physics, 115(4):1716–1733, 2001.

[55] Matt J Keeling and Pejman Rohani. Modeling infectious diseases in humans and animals. Princeton university press, 2011.

[56] Stefan Sellman, Kimberly Tsao, Michael J Tildesley, Peter Brommesson, Colleen T Webb, Uno Wennergren, Matt J Keeling, and Tom Lindström. Need for speed: An optimized gridding approach for spatially explicit disease simulations. PLoS computational biology, 14(4):e1006086, 2018.

[57] Micah Allen, Davide Poggiali, Kirstie Whitaker, Tom Rhys Marshall, Jordy van Langen, and Rogier A Kievit. Raincloud plots: a multi-platform tool for robust data visualization. Wellcome open research, 4:63, 2021.

[58] IShin Tseng, Bing-Yi Pan, Yen-Chen Feng, and Chi-Tai Fang. Re-evaluating efficacy of vaccines against highly pathogenic avian influenza virus in poultry: a systematic review and meta-analysis. One Health, page 100714, 2024.

[59] Carla Correia-Gomes and Nick Sparks. Exploring the attitudes of backyard poultry keepers to health and biosecurity. Preventive veterinary medicine, 174:104812, 2020.

