## Supplementary Information for "A modelling assessment for the impact of control measures on highly pathogenic avian influenza transmission in poultry in Great Britain"

#### Data

The demographic data on poultry premises includes the centroid of each premises polygon (defined as a CPH — County/Parish/Holding number — entity). The poultry case data provides a latitude and longitude value of the case. To match the correct premises from the demographic data to the reported IP in the case data, we found the closest premises in the demographic data to the case coordinates with similar poultry populations between the two data sets. Specifically, we consider all premises within a 2 km radius of the case data coordinates and match the closest premises with the same bird species types and the same size of premises in terms of number of birds (less than 50, from 50 to 1,000 or more than 1,000). If no match is found to all the criteria, we match the remaining premises if there is a valid premises in the demography data in very close proximity (less than 200 m) or if the bird type does not match, but the numbers of birds do match, and vice versa. Premises with still no match are added to the demographic data set. Of the 200 premises in our data set, 158 were matched in the first step, a further 6 were matched by close proximity, and another 6 were matched by number of birds or bird type. The final 30 premises were added to the data set.

Our distribution of poultry premises sizes is given in a histogram (Figure S1A). There are many small-sized premises (less than 100 birds) with a decreasing number of premises with larger sizes. However, there is a second small peak of premises with a large number of birds (approximately 20,000 birds). Spatially, most poultry is kept in rural areas of Great Britain with a relatively high density in the East of England, South West and close to the border with Wales in the West Midlands (Figure S1B). The most common poultry type kept are Galliformes (Figure S1C) with approximately 282 million, compared to 4 million waterfowl and 43 million other birds. The most common premises type is Galliformes only, followed by mixed, other birds only, and then waterfowl only, with many fewer premises (Figure S1L). The distributions of birds in premises of particular types shows that, similarly to the total number of birds, Galliformes only and waterfowl only premises are bimodal in size, with birds kept in small or large numbers with two peaks in the distribution (Figure S1D and E). Other birds only premises commonly have moderate numbers of kept birds (Figure S1F), and mixed species type premises have a similar distribution to all premises types combined (Figure S1G). Spatial distributions of total bird species type numbers are shown in Figure S1I–K.

For considering spatial strategies, when we use enhanced biosecurity in the main text, we use different spatial scales, including at the county and region levels. These are all areas of different shapes and sizes and are displayed in Figure S2. There are 207 counties included and 11 regions in Great Britain.

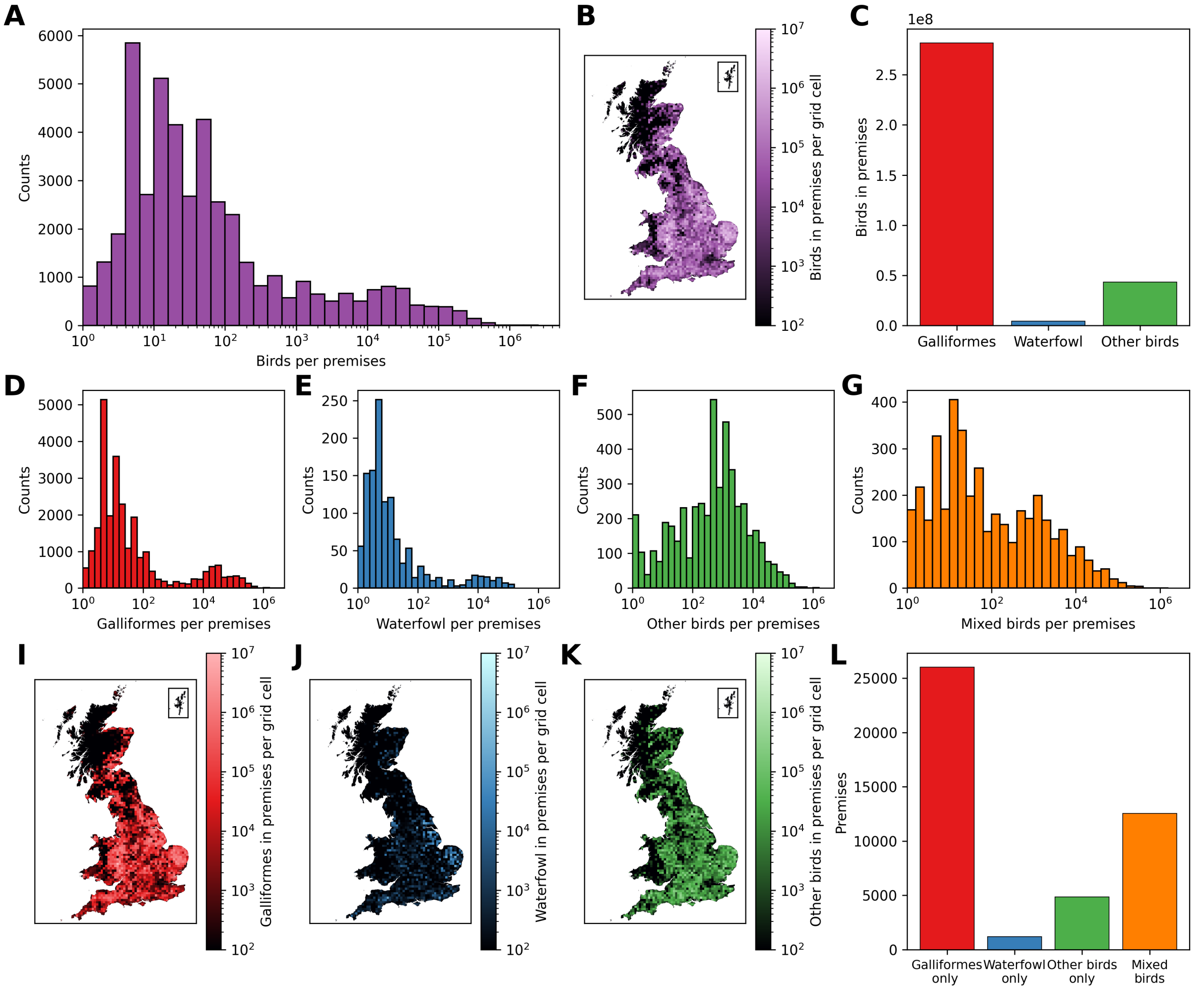


Figure S1: **Summary of poultry premises data.** (A) Distribution of the total number of birds on each poultry premises in the data set. The x-axis is shown on a logarithmic scale. (B) Spatial distribution for the total number of birds within all premises within a 10 km by 10 km grid cell. (C) The total number of birds for each bird type across all poultry premises. (D–G) Distribution of the number of birds divided by premises with: (D) Galliformes only, (E) waterfowl only, (F) other birds only and (G) mixed species premises. A logarithmic scale is used for the x-axis. (I–K) Spatial distribution for the total number of birds of each species within premises within a 10 km by 10 km grid cell for: (I) Galliformes, (J) waterfowl and (K) other birds. (L) Number of premises for each species type: Galliformes only, waterfowl only, other birds only and mixed birds.

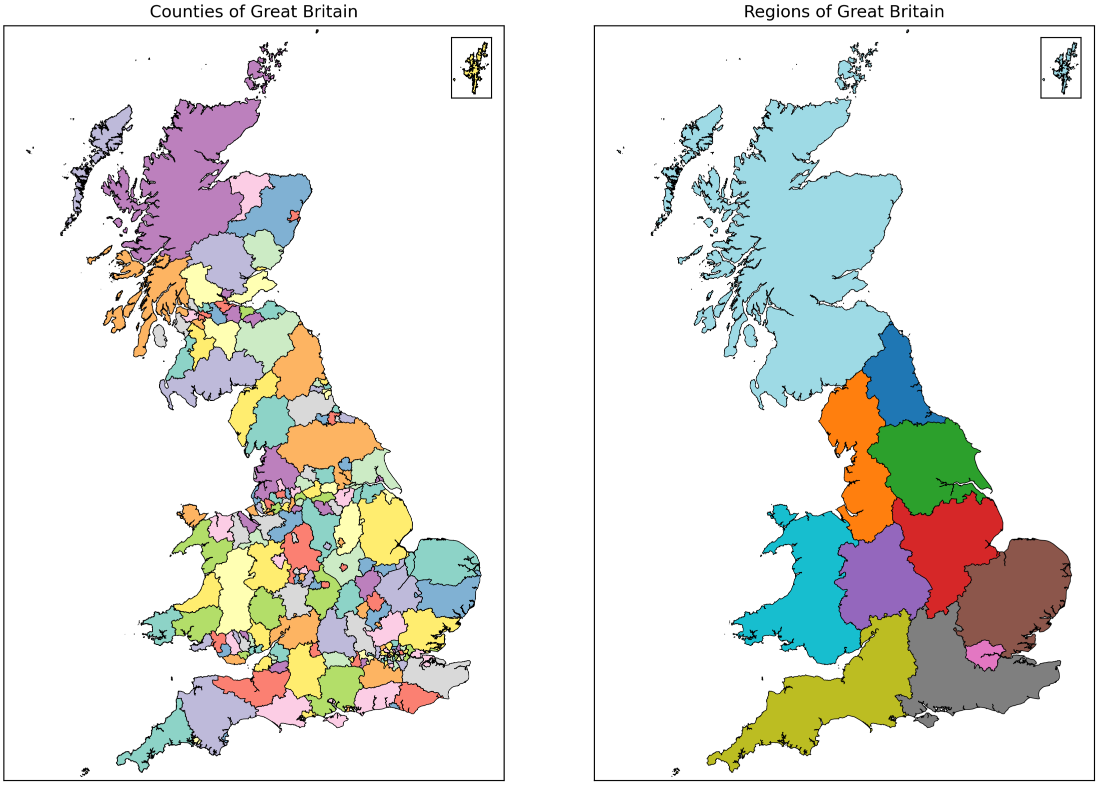


Figure S2: **Counties and regions of Great Britain.** Boundaries of the counties and regions are shown on a map. Source: Office for National Statistics licensed under the Open Government Licence v.3.0. The shapefiles used can be found at `https://www.arcgis.com/home/item.html?id=2cc79b94571a438bb05bc9ec16831529` and `https://geoportal.statistics.gov.uk/datasets/5a393192a58a4e50baf87eb4d64ca828_0/explore` respectively.

#### Model fitting

We assume the infection events occurred according to a continuous-time non-homogeneous Poisson process, given infection rate $\lambda_j(t)$. We use this method for computational efficiency in exploring the state space, noting that the approximation from continuous time inference to discrete time simulation will provide negligible differences. The time to notification from being infectious $(N_i - I_i)$, is also assumed to be gamma distributed. Therefore, the joint posterior of the parameters can be described by:

$$\pi(\theta \mid N) \propto \left(\prod_{i=1}^{n_I} \lambda_i(E_i)\right)\left(\exp\left(-\sum_{t=1}^{T}\sum_{i=1}^{n} \lambda_i(t)\right)\right)\left(\prod_{i=1}^{n_I} (N_i - I_i)^{a-1} \exp\left(-b\left(N_i - I_i\right)\right)\right)\left(\prod_{p=1}^{|\theta|} f_p(\theta_p)\right), \quad (1)$$

where $n$ is the total number of premises and $n_I$ is the total number of these premises that are infected up to time $T$. The prior distributions of the fitted model parameters are given by $f_p((\theta_p)$ for parameters $\theta_p$, where $\boldsymbol{\theta} = \{\epsilon_0, \gamma_0, \gamma_1, \delta, \psi_0, \psi_1, \psi_2, \phi_0, \phi_1, \phi_2, \xi_1, \xi_2, \zeta_1, \zeta_2, \nu_0, \nu_1\}$. In practice, we fit log-transformed parameters and account for this in the MCMC accordingly.

We use a reversible Markov chain Monte Carlo (MCMC) method, which proposes a new set of parameters in each MCMC iteration and then proposes a fixed number of additional events for uniformly chosen random premises, which are:

i) Update the time between a premises becoming infectious and being notified,

ii) Add an occult infection as a new infection that has not yet been detected at the end of the simulation period,

iii) Remove a previously added occult infection.

We choose the fixed number of additional events to be equal to 5% of the total number of infected premises.

We perform 211,000 MCMC iterations for two independent chains, which for the first 10,000 we update each parameter separately in order to get a better initial starting point before updating all parameter values as a set simultaneously for the following iterations (Figure S3A). For obtaining posterior distributions, we remove these first 10,000 iterations and an additional 1,000 of the subsequent iterations as a burn-in period, leaving 200,000 iterations from which we obtain posterior estimates.

These chains visually appear to have converged (Figure S3A), however, we also calculate the convergence of these chains using the Gelman–Rubin statistic [1] (Figure S3B). This confirms that as we increase the number of MCMC iterations, the Gelman–Rubin statistic trends towards a value of 1, and so we have likely performed a sufficient number of iterations.

For the posterior parameter sets, we take the 200,000 MCMC iterations and thin these to reduce the auto-correlation and get a set of 1,000 parameter sets for each chain for a total of 2,000 posterior parameter sets. We observe that none of these parameters diverge significantly for the prior estimates, but do form smooth distributions that are unique from the priors, indicating that the fitting process has been successful (Figure S4).

Similarly, we output a unique time to notification posterior distribution for each of the infected premises. We show an example of three premises of the 200, which all exhibit similar features, alongside the distribution of the average time to notification time across all the premises (Figure S5). We see that during the fitting process, the algorithm is generally selecting for a shorter time to notification than given by the priors, with a mean value of approximately 6.7 days before notification.

The number of occult infections remains low, with the posterior only including between zero and ten additional premises, with two being the most likely value (Figure S6). This is to be expected since the end of the simulations is at the end of the summer (30 September 2023), where infections are typically at a low number before migratory wild birds arrive in Great Britain. We show the probability that each premises is included as an occult infection, where the largest probability is 0.0012, demonstrating that all premises are much more likely to be not infected. The premises with the highest probability of being occult is the closest premises to the last recorded infection (on 26 September 2023), indicating that the model results are consistent with the data. Indeed, we truly observed zero IPs for the time scale of occult infections on 30 September 2023, in agreement with the modelled results.

Marked on the map in Figure S6 are red crosses and dates, which correspond to the infected premises in the last month of the data period. There are no premises with large probabilities close to the infected premises

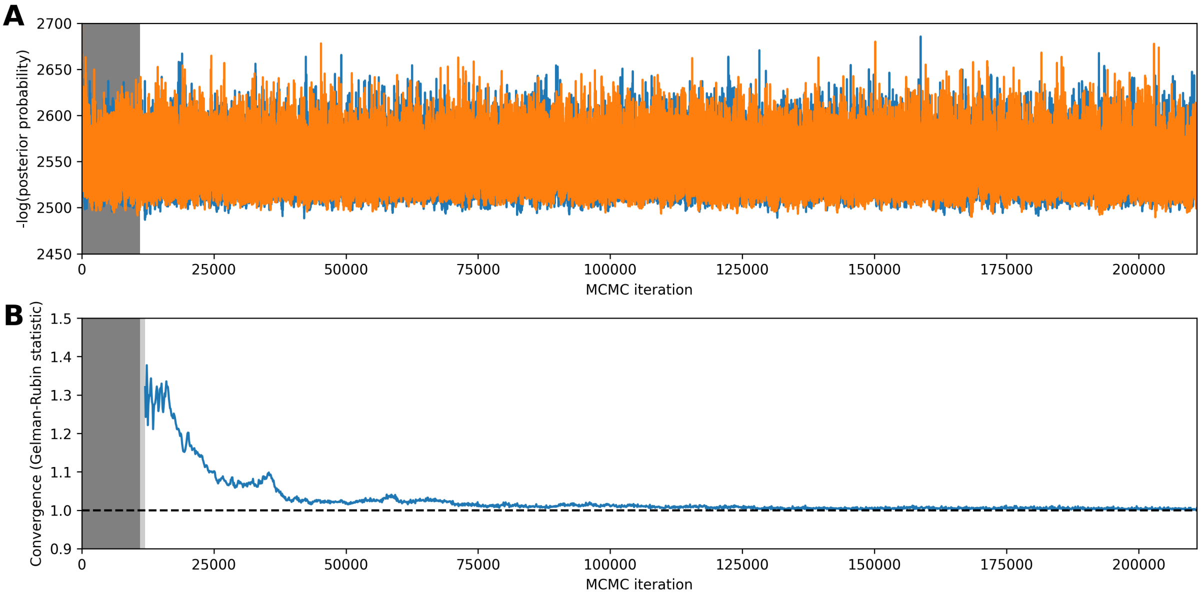


Figure S3: **MCMC chain and convergence.** (A) The two MCMC chains showing the negative logarithm of the posterior probability by iteration. A burn-in of 11,000 iterations is used (shown by the dark grey background), followed by a further 200,000 iterations. (B) Convergence of the two chains by MCMC iteration as calculated by using the Gelman-Rubin statistic on 1000 thinned samples. The dashed line at 1 shows the chains converging towards this value as the MCMC process is simulated. The dark grey region shows the neglected burn-in period, and the lighter grey region shows the period before 1,000 values post-burn-in are available to calculate the statistic.

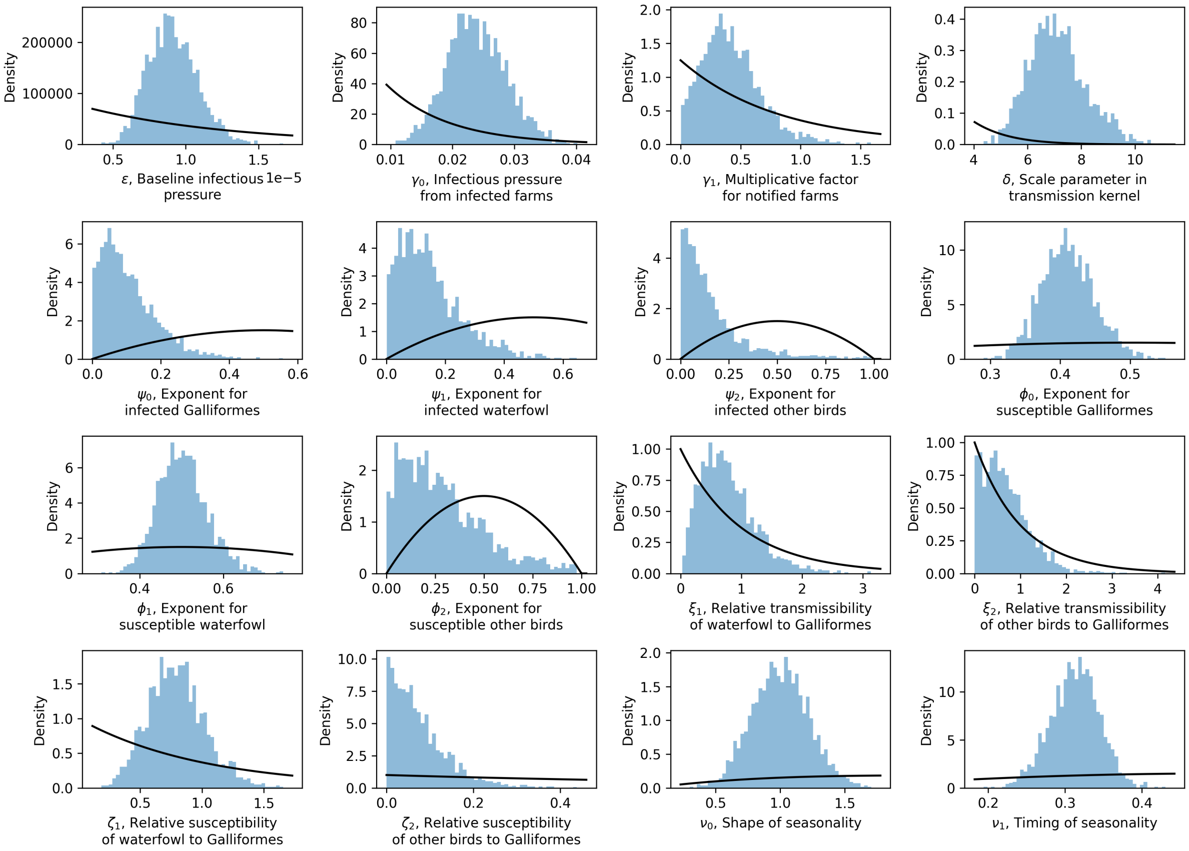


Figure S4: **Posterior parameter distributions.** Parameter distributions for the 16 model parameters after Markov chain Monte Carlo. The black lines show the prior distributions used.

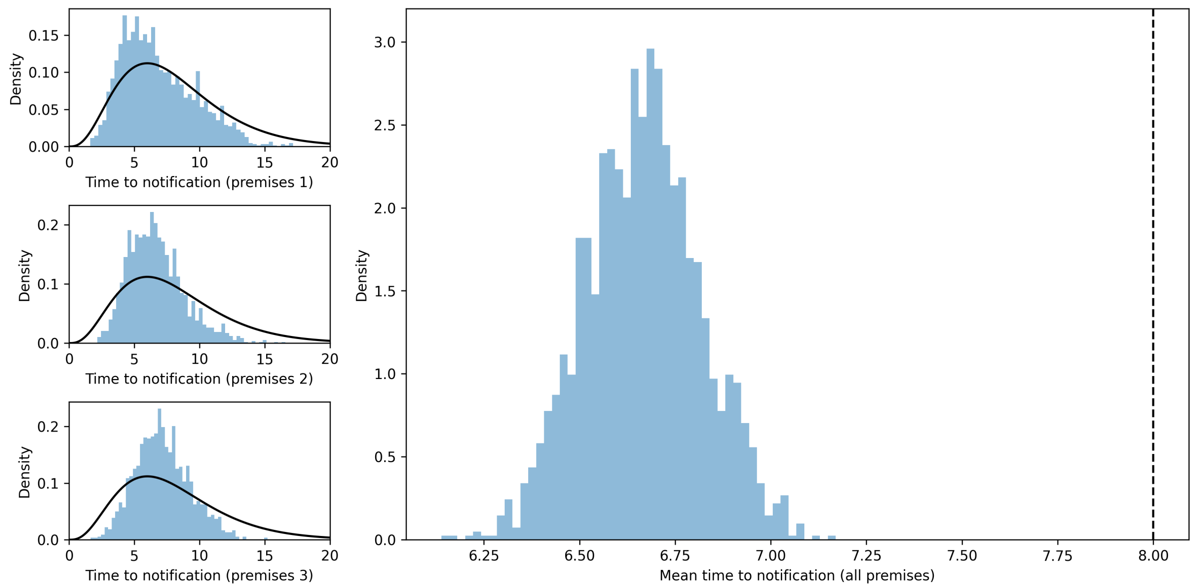


Figure S5: **Distributions of the time to notification.** Time to notification distributions for three example premises of the 200 infected premises in the 2022–23 season. The black lines show the prior distribution, which is the same for each premises. To the right is the distribution of the mean value across the 200 infected premises, where the dashed line is the mean of the prior distribution.

other than the premises infected on 26 September, as they are not sufficiently close to the end of the modelled period compared to the incubation period plus the time to notification. The other premises with relatively large probabilities of being occult infections (albeit still less than 0.001) are typically premises with large numbers of kept birds.

### Model assumptions

Throughout the main text, we assumed the transmission kernel takes a Cauchy form (Main Text, Equation 4). Here, we also considered an exponential form given by:

$$K_{ij} = K\left(d_{ij}\right) = A \exp\left(-\delta d_{ij}\right), \tag{2}$$

where A is a constant such that the median value of $\gamma_0$ remains unchanged. The parameter distributions after MCMC are shown in Figure S7.

The posterior parameter estimates remain broadly unchanged, with the exception of the $\delta$ parameter, which is shown in separate panels due to different scales. This is different because the parameterisation is inherently different due to the differing functional form of the kernel. However, the overall shape of the transmission kernels are similar (Figure S8).

The mean exponential kernel is smaller at close distances, is larger for moderate distances and smaller again for very large distances. This is despite there being a strong overlap in the uncertainty. Due to the minimal difference, we chose to use the Cauchy form of the kernel in the main text because of the higher transmission at closer distances.

Seasonality is included in the model within the background infection term:

$$\epsilon(t) = \epsilon_0 \exp\left(-\nu_0\left(1 + \cos\left(2\pi\left(\frac{t}{365} - \nu_1\right)\right)\right)\right), \tag{3}$$

In the fitted model, we see that the posterior parameter distributions give a seasonality relatively sharply across November to February and is likely to reach its maximum in late December. Through much of the spring and summer, transmission is much reduced with the seasonality factor close to 0.2 (Figure S9).

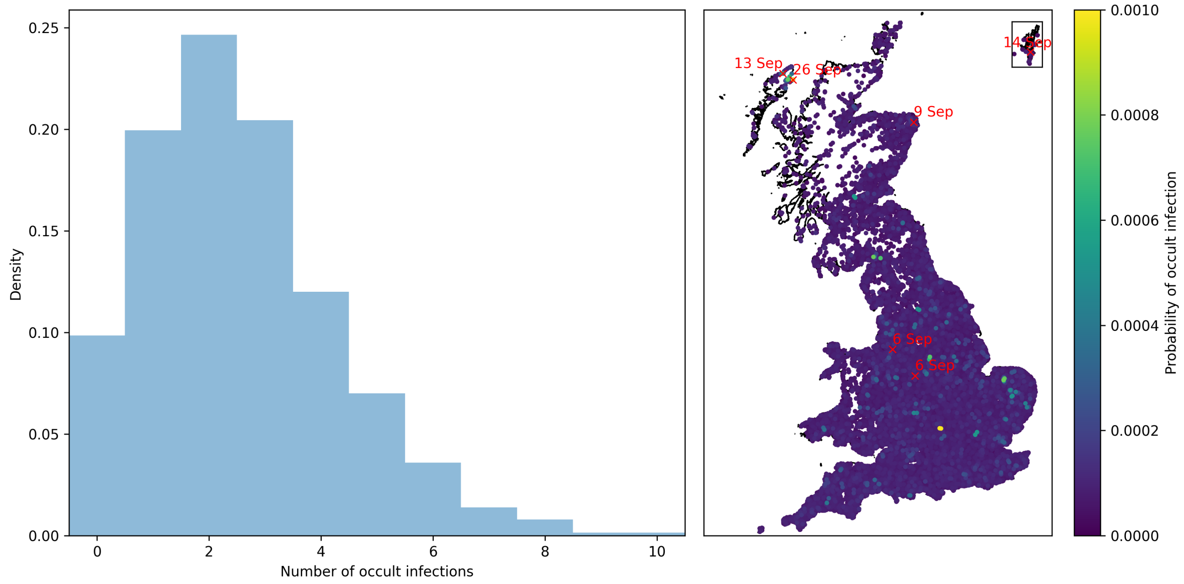


Figure S6: **Occult infections.** Distribution for the number of occult infections and a map of the probability that each premises is an occult infection. Additionally marked in red are the true infected premises in the last month of the season, with dates of notification shown.

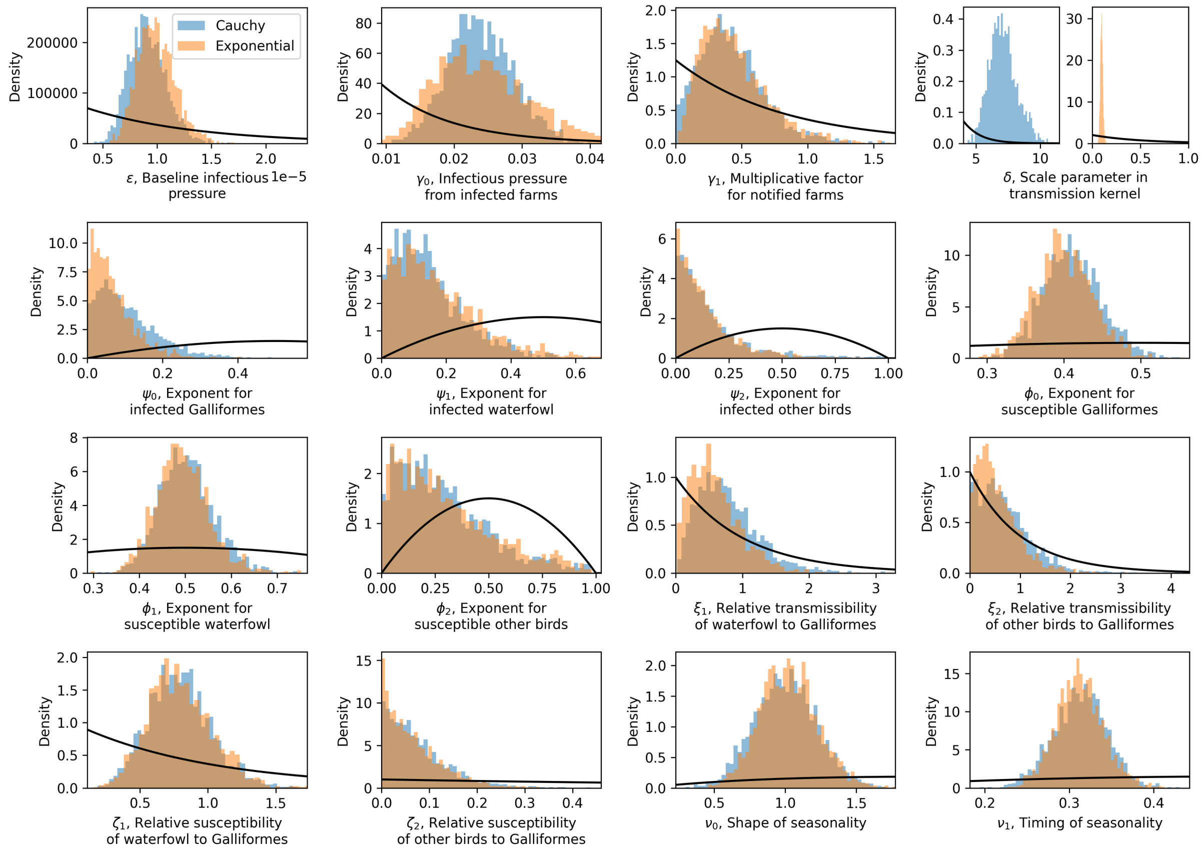


Figure S7: **Posterior parameter distributions for models with Cauchy and exponential transmission kernels.** Parameter distributions for the 16 model parameters after Markov chain Monte Carlo. The Cauchy kernel of the main text is shown in blue, and the exponential kernel is shown in orange. The black lines show the prior distributions used.

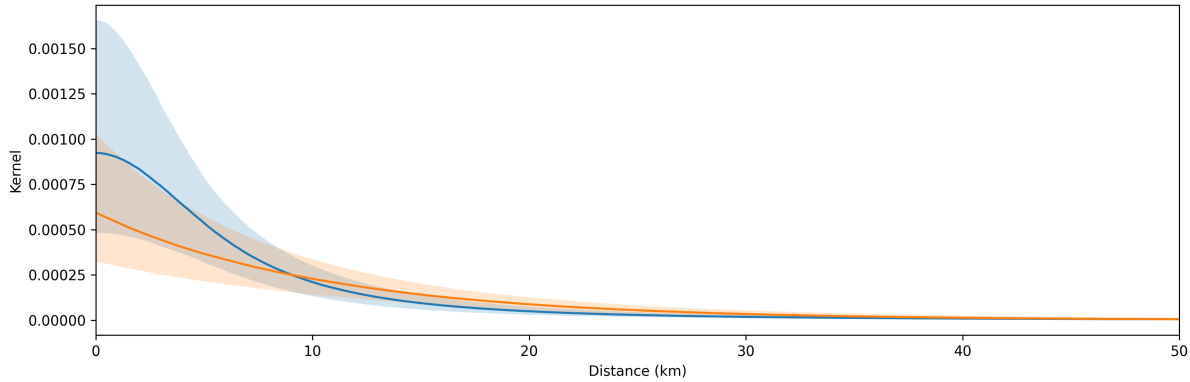


Figure S8: **Transmission kernels.** The fitted functional form of the transmission kernels is shown with 95% prediction intervals using the posterior parameter distributions. The Cauchy kernel of the main text is shown in blue, and the exponential kernel is shown in orange.

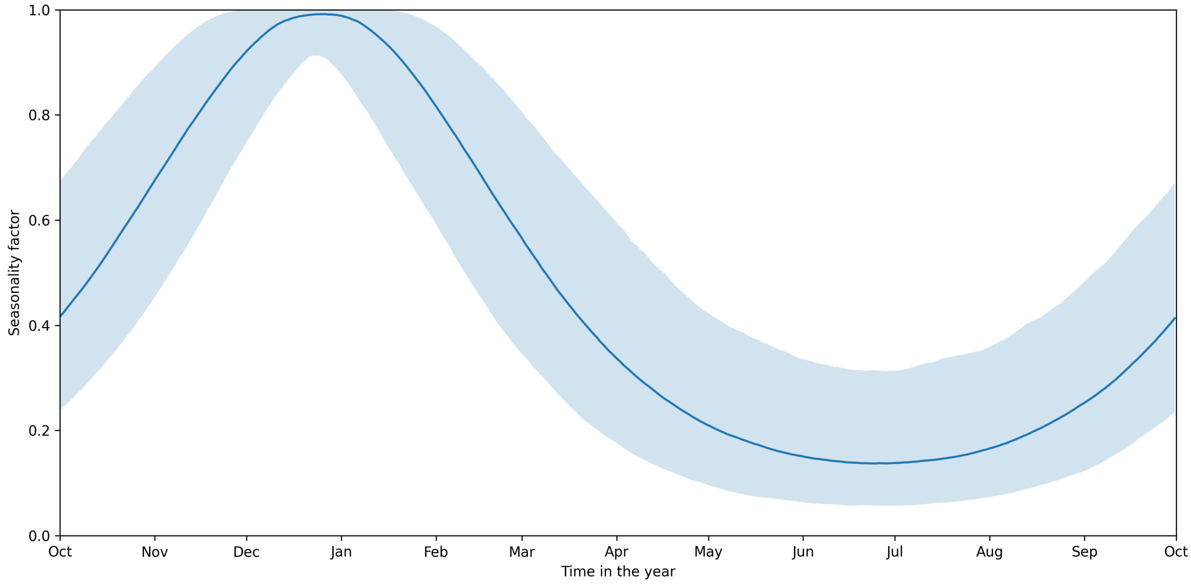


Figure S9: **Seasonality of background infections.** The fitted functional form of the seasonality factor is shown with 95% prediction intervals using the posterior parameter distributions. The seasonality factor is an equation that multiplies with the fitted parameter $\epsilon_0$ to give $\epsilon(t)$.

#### References

[1] Andrew Gelman and Donald B Rubin. Inference from iterative simulation using multiple sequences. *Statistical science*, 7(4):457–472, 1992.
